# Automation Assisted Anaerobic Phenotyping For Metabolic Engineering

**DOI:** 10.1101/2021.05.03.442526

**Authors:** Kaushik Raj, Naveen Venayak, Patrick Diep, Sai Akhil Golla, Alexander F. Yakunin, Radhakrishnan Mahadevan

## Abstract

Microorganisms can be metabolically engineered to produce a wide range of commercially important chemicals. Advancements in computational strategies for strain design and synthetic biological techniques to construct the designed strains have facilitated the generation of large libraries of potential candidates for chemical production. Consequently, there is a need for a high-throughput, laboratory scale techniques to characterize and screen these candidates to select strains for further investigation in large scale fermentation processes. Several small-scale fermentation techniques, in conjunction with laboratory automation have enhanced the throughput of enzyme and strain phenotyping experiments. However, such high throughput experimentation typically entails large operational costs and generate massive amounts of laboratory plastic waste. In this work, we develop an eco-friendly automation workflow that effectively calibrates and decontaminates fixed-tip liquid handling systems to reduce tip waste. We also investigate inexpensive methods to establish anaerobic conditions in microplates for high-throughput anaerobic phenotyping. To validate our phenotyping platform, we perform two case studies - an anaerobic enzyme screen, and a microbial phenotypic screen. We used our automation platform to investigate conditions under which several strains of *E. coli* exhibit the same phenotypes in 0.5 L bioreactors and in our scaled-down fermentation platform. Further, we propose the use of dimensionality reduction through t-distributed stochastic neighbours embedding in conjunction with our phenotyping platform to serve as an effective scale-down model for bioreactor phenotypes. By integrating an in-house data-analysis pipeline, we were able to accelerate the ‘test’ phase of the design-build-test-learn cycle of metabolic engineering.

## Introduction

Microbial production of chemicals has gained prominence in the past few decades due to rising populations and increased concerns over the sustainability of conventional means of chemical production. Advances in metabolic engineering and synthetic biology have enabled the generation of mutant strains that are adept at producing a wide range of natural and non-natural chemicals^32^. However, a myriad of scale-up issues can arise at increasingly larger scales, that could render many microbial production platforms economically infeasible^8,24^. Hence, several iterations of the design-build-test-learn (DBTL) cycle (Figure 1a) may be required at smaller scales before moving on to production in larger scale bioreactors.

**Figure 1:**
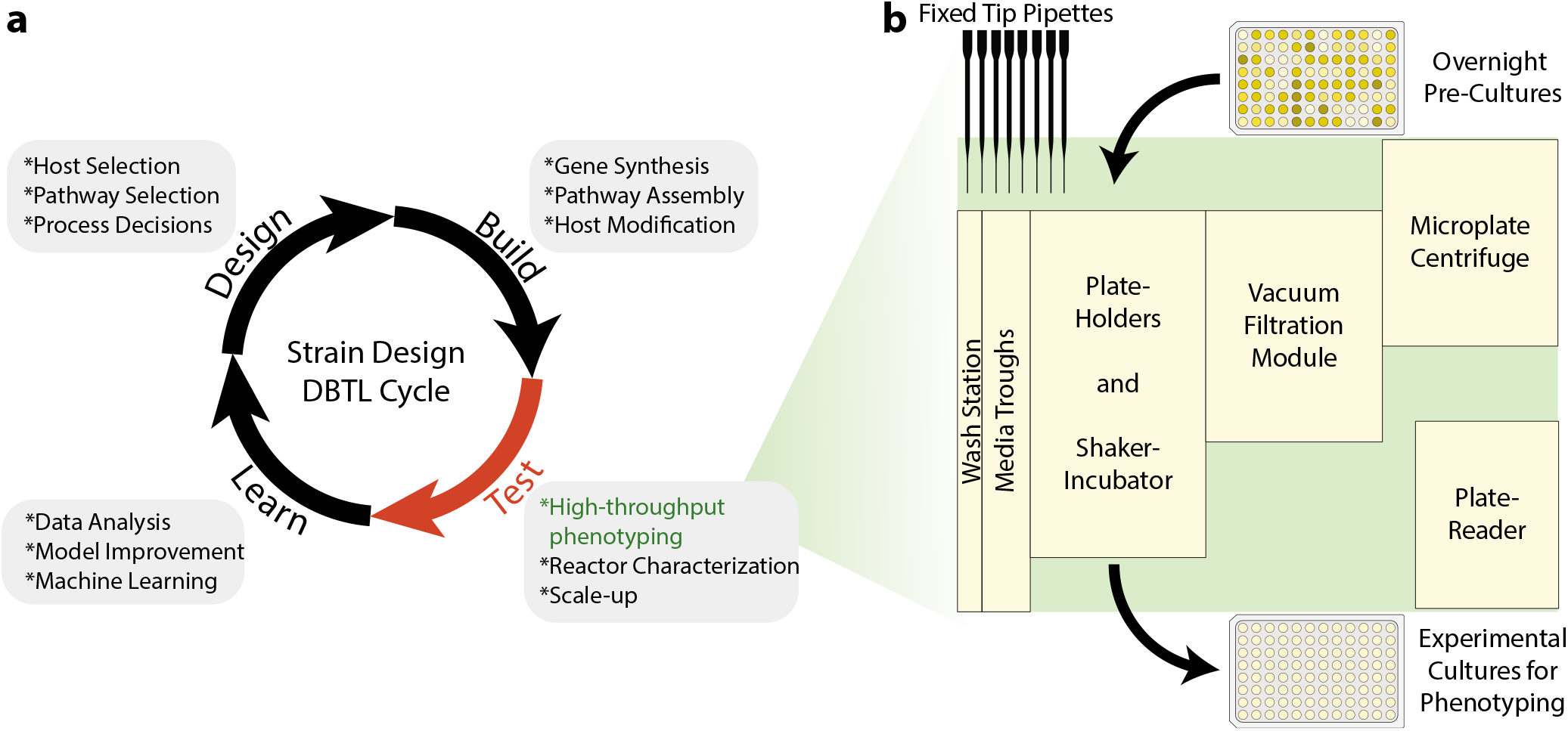
DBTL cycle in metabolic engineering and layout of phenotyping platform. **a.** Typical decisions and tasks involved in each step of the DBTL cycle for strain engineering. Test cycle remains a bottleneck due to the time and costs incurred in phenotyping a large number of strains/enzymes. **b.** Deck layout of liquid handling platform used in this study. Relatively few equipment can be assembled and repurposed to establish an effective high-throughput phenotyping platform.

The development of genome-scale metabolic models and computational tools that use these models to predict genetic interventions for strain design has assisted the ‘design’ phase of the DBTL cycle^4,11,43,52^. Similarly, advances in DNA synthesis, computational tools to streamline DNA assembly, and the establishment of DNA foundries around the world have also allowed for the rapid construction of mutant strain and enzyme libraries that incorporate these intervention strategies, accelerating the ‘build’ phase^5,16,18,27,37^. The ‘test’ phase i.e. characterization/phenotyping of the strain and enzyme libraries generated in the ‘design’ and ‘build’ phases of the DBTL cycle remains a key bottleneck. The prohibitive cost of analyzing the phenotypes of all microbial strains in the generated mutant libraries using laboratory scale bioreactors necessitates the development of standardized high-throughput, small-scale protocols to characterize them. Recently, several machine learning techniques have been adapted for metabolic engineering applications, with several tools being developed that promise to assist the ‘learn’ phase^31,41^. These tools also necessitate the generation of large and reliable experimental phenotypic datasets that are only economically feasible at extremely small scales, further bolstering the need for protocols for high-throughput phenotyping platforms ^6^.

In the recent past, there have been several attempts to develop small scale fermentation platforms using miniature bioreactors and specialized microplates to cultivate and characterize strains, increasing experimental throughput^2,25,29^. However, the operational costs of using such systems is quite high due to the requirement of specialized microplates and intricate pH control mechanisms. Further, the automation of strain cultivation and other routine workflows to enhance throughput using such systems may be very expensive to implement. The earliest attempts at high-throughput fermentation were through the use of standard 96-well microtiter plates for parallel cultivation of microbes.^10^. The low cost and enhanced throughput of these systems made them very valuable to perform preliminary screens on a large number of strains. However, these systems suffer from several disadvantages including increased rates of sample evaporation and reduced oxygen transfer. Therefore, microbial phenotypes observed in these scales may not be replicable at the scale of bench-top reactors under aerobic conditions. Yet, these systems may still be suitable to phenotype microbes under anaerobic conditions where oxygen transfer is not crucial. *E. coli* can be engineered to produce an array of commercially important compounds such as lactic acid under anaerobic conditions^9,35^. Moreover, the production phase of many industrial fermentation processes involve high density cultures where oxygen transfer is limited. Microtiter plates are particularly suited for anaerobic fermentations due to the inherent difficulty in achieving high oxygen transfer rates and have the potential to be able to replicate the phenotypes of microbes observed in bench-top bioreactors.

The advent of liquid handling systems has assisted in the use of such small-scale fermentation platforms, enhancing throughput by reducing human effort and time required to set up phenotyping experiments^7,17,44,47^. Use of such automation systems also enhances the reproducibility of experiments through the use of standardized protocols. While automated liquid handling platforms can rapidly accelerate the throughput of experiments, maintaining sterile conditions during long high throughput workflows is challenging. Contamination arising from the environment can be effectively curbed through the use of HEPA filters^20^. However, cross-contamination resulting from tip carryover could still be a problem, since any residual contaminant in the components of the platform could potentially confound results from a large set of experiments. Liquid handling systems with disposable tips have been successfully adapted to cultivate cells and perform other routine microbiological workflows with minimal contamination^20,26,46^. These systems simply discard used and contaminated tips after each pipetting step, thereby eliminating contamination. This would inevitably result in massive amounts of plastic waste when such systems are used for high-throughput workflows. The rapidly increasing adoption of automated workflows in research laboratories would only exacerbate this problem due to their increased throughput^19^. Moreover, the need for a massive number of sterilized tips would increase the operational costs required to implement such workflows^28,46^. The use of fixed-tip liquid handlers with effective decontamination protocols could address concerns about sustainability and operational costs.

In this work, we describe several efforts towards enhancing the utility of fixed-tip liquid handling systems for automated high-throughput phenotyping using a platform consisting of a fixed-tip liquid handler, microplate centrifuge, plate-reader, vacuum filtration module, plate handling robot, and a shaker incubator (Figure 1b). To this end, we develop decontamination protocols to eliminate microbial carry-over and cross-contamination in fixed-tip liquid handlers, describe an automated calibration workflow to calibrate liquid handling pipettes, and establish relatively easy methods to ensure anaerobicity of media for anaerobic phenotyping. Then, we validate our platform by performing an anaerobic enzyme screen and investigate conditions that allow reasonable replication of bioreactor microbial phenotypes in 96-well microplates.

## Results & Discussion

### A decontamination protocol for fixed-tip liquid handlers

Fixed-tip liquid handling systems require decontamination after every pipetting step to curb biological cross-contamination. A disinfection step where tips are washed and incubated with ethanol has been proposed in the past to address contamination issues^44^. However, this protocol required the incubation of pipette tips in ethanol for five minutes between each pipetting step, reducing the throughput of this system. More recently, one study used a layer of ethanol, aspirated immediately before aspirating biological samples to maintain sterility.^23^. While this protocol is faster, it may result in reduced cell viability due to direct contact between the disinfectant and cells.

To address these issues, we examined the effectiveness of a simple decontamination protocol that uses a solution of sodium hypochlorite (bleach) to disinfect pipette tips (Figure 2a). In order to simulate typical contamination events during cell culture workflows, we programmed the pipette to aspirate 200 μL of viable *E. coli* cells in their exponential phase of growth, hold for 30 seconds with the pipette tips dipped inside the culture, and dispense the cells back into the solution. Then, the tips aspirate 400 μL of bleach, hold for a specified interval - *’t’* seconds with the tips dipped inside, and dispense the disinfectant. We repeat this bleach wash for a specified number of times - *’n’* and when complete, wash the tips with system liquid - sterilized ultrapure water, to remove any traces of the disinfectant. Finally, to examine the effectiveness of our decontamination procedure, we aspirate 200 μL of sterile LB media from a microplate, hold for 30 seconds and dispense back into the same wells. Any persisting *E. coli* cells in the tips would lead to contamination of the media and show cell growth after incubation of the plate. We used a wash with water as a negative decontamination control to ensure that contamination events are captured effectively using this procedure.

**Figure 2:**
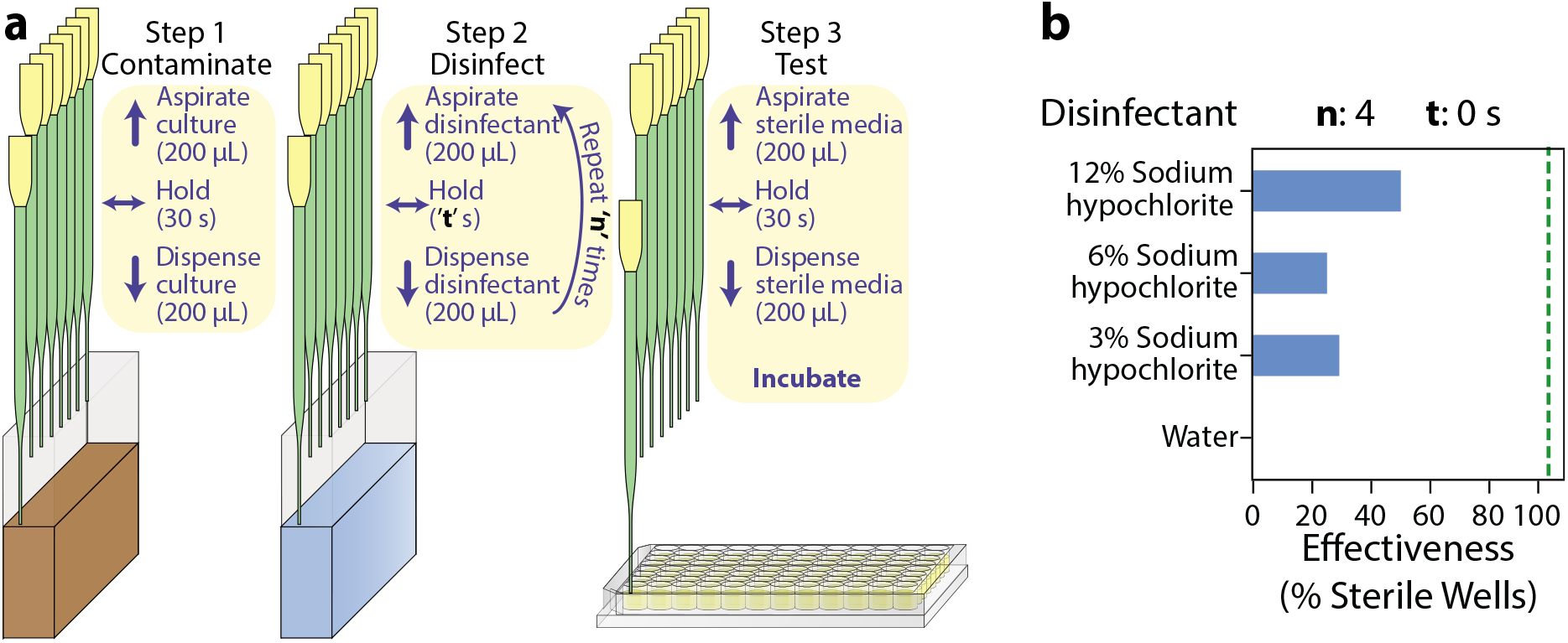
Preliminary decontamination protocol. **a.** Steps to decontaminate and investigate effectiveness of the decontamination protocol. **’n’** represents the number of washes with the disinfectant and **’t’** represents the duration for which the disinfectant is held within the tips for each wash. **b.** Initial decontamination test using different concentrations of sodium hypochlorite(bleach) with **’n’**=4 and **’t’**=0 and the default air-gap of the system (10 μL). Each bar represents effectiveness calculated from 24 replicates.

First, we examined the efficacy of this procedure using varying concentrations of bleach, with *’n’*=4 washes and zero hold time (’*t*’ = 0 s). The sterilization effectiveness was calculated as the percentage of contaminated wells resulting from the corresponding decontamination procedure. As seen in Figure 2b, the negative control - water resulted in zero effectiveness. Increasing the concentration of bleach seemed to positively impact the effectiveness of our protocol. However, even at the highest concentration of bleach, we only observed a 50% effectiveness of decontamination. We considered that varying the number of washes - *’n’* and the hold time for the disinfectant - *’t’* could improve our system due to longer contact with bleach. Increasing the number of washes and the hold time indeed had a positive impact on the sterilization effectiveness, with the best values being achieved at the highest levels of *’n’* and *’t’* (Figure 2d - top-left panel). However, this was still unacceptable as the target was to completely eliminate contamination events. Moreover, operating at the highest levels of *’n’* and *’t’* increased the run-time of the decontamination protocol to about 1 minute and would therefore reduce the throughput of our system.

Upon further investigation of the pipetting protocol, we observed that like most fixed-tip liquid handling systems, our pipettes aspirate a very small amount of air (10 μL) before each pipetting step to separate the system liquid from the liquid being pipetted - the process liquid(Figure 3a). By increasing this air-gap, we were able to remarkably improve our decontamination protocol, achieving complete sterilization using an air-gap of 250 μL (Figure 3b and Supplementary Figure S1). Interestingly, at the highest level of air-gap, we observed zero contamination events even at our lowest levels of *’n’* and *’t’*. It appears that when the volume of the air-gap is less than the maximum operating volume of the process liquid, there is a possibility for the sterile system liquid to come in direct contact with parts of the pipette that have not yet been disinfected. The system liquid is therefore compromised and could harbour viable cells, which increases the possibility of contamination during further pipetting steps (Figure 3c). An air-gap greater than the highest process volume ensures complete separation of the system and process liquids, leading to proper decontamination(Figure 3c). We found that our protocol remained effective over a range of bleach concentrations and with 70% ethanol even at the lowest levels of *’n’* and *’t’*((Figure 3d). For all further experiments, we chose to use two washes with 6% bleach as the disinfectant. The duration of the entire decontamination procedure is about 10 seconds and is therefore at par with the throughput achieved using disposable plastic tips, with no plastic waste generated and minimal amounts of disinfectant used.

**Figure 3:**
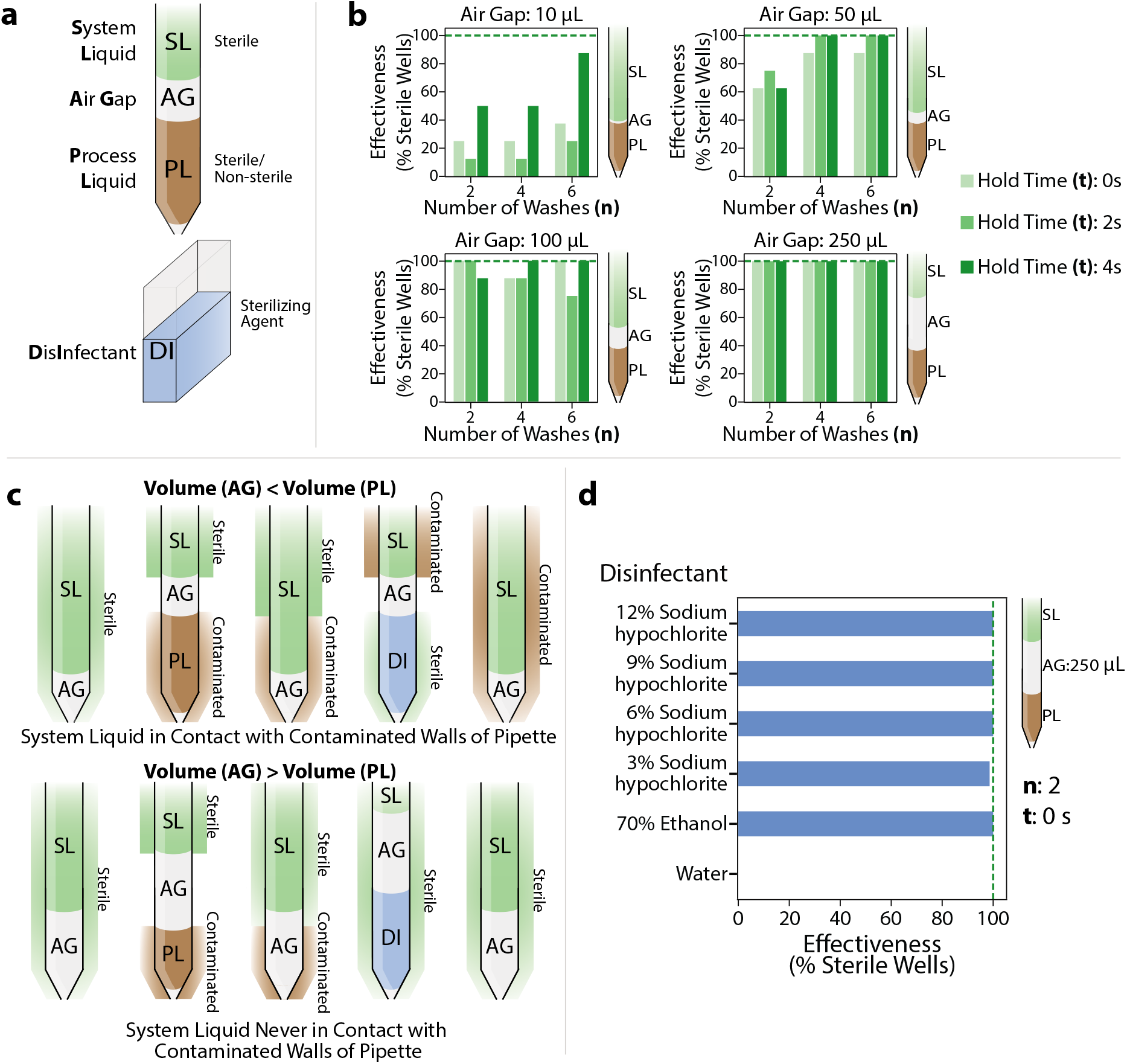
Optimizing decontamination protocol. **a.** Schematic showing tip layout during a typical pipetting step. **b.** Effect of varying the air-gap on the effectiveness of sterilization using 12% sodium hypochlorite for different values of **’n’** - number of disinfectant washes and **’t’** - disinfectant hold time. Each bar represents effectiveness calculated from 8 replicates. Negative controls using water as the disinfectant resulted in zero sterilization effectiveness for all values of **’n’** and **’t’**. **c.** Proposed mechanism for enhanced sterility upon increasing the volume of the air-gap to be larger than pipetted volumes. **d.** Sterilization effectiveness for different disinfectants with an air-gap of 250 μL. Each bar represents effectiveness calculated from 72 replicates.

### Automated photometric calibration of liquid handling pipettes

Following the implementation of our decontamination protocol, we observed that the accuracy of the pipettes had diminished quite significantly, with aberrant volumes being pipetted consistently. In order to examine the pipetting accuracy of the liquid handler before and after changing the air-gap, we used a photometric assay to compare the volumes pipetted by the automated platform to manually pipetted standards, similar to an assay described previously^45^. In our assay, we used an aqueous solution of potassium dichromate (K_2_Cr_2_O_7_) within concentration ranges that showed a linear relationship with absorbance at 350 nm, as a photometric standard. We pipetted different levels of the standard within volume ranges required during routine operation (3 μL - 200 μL) into a microplate. Then, an on-deck plate reader was used to measure the absorbance and determine the concentration of samples in each well, thereby providing an accurate estimate of the pipetted volumes. We observed that after increasing the air-gap, the pipetting error increased significantly for all pipette tips (Figure 4a), with values of up to 40% for some tips, implying that pipetting accuracy would depend on the volume of air-gap used for each pipetting step. The deviations in pipetted volumes were well above the maximum acceptable limits specified by the International Organization for Standardization^22^ and would certainly hinder normal operation of the platform.

**Figure 4:**
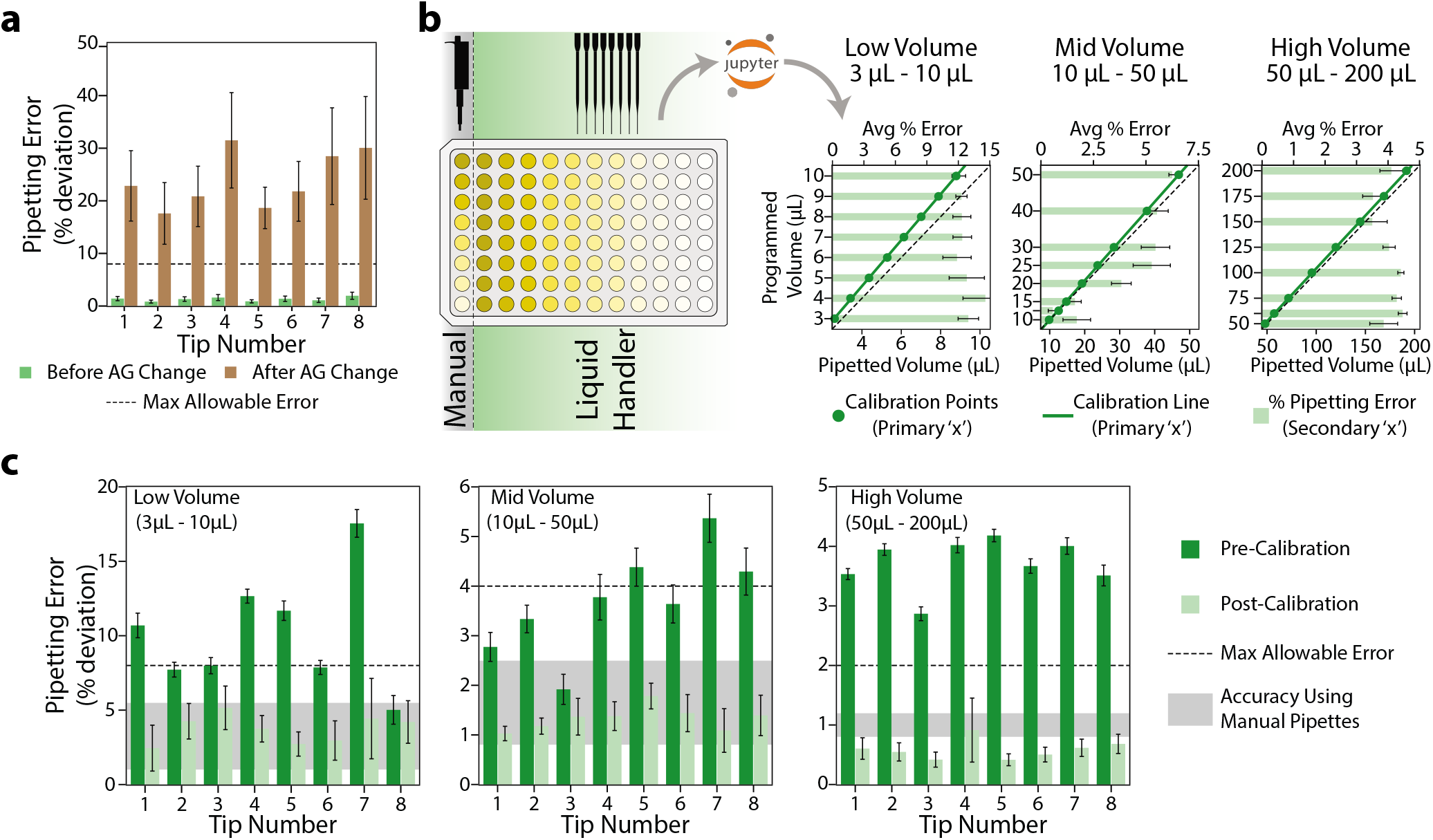
Automated Photometric Pipette Calibration. **a.** Change in pipetting error due to an increase in the air gap. **b.** Workflow for automated photometric calibration. The liquid handler is made to pipette a photometric standard at different levels onto a microplate. The absorbance data of the microplate are recorded and fed to a pythonic script which automatically calculates pipetting errors and calibration parameters for the pipette. **c.** Pre and post-calibration pipetting error with the airgap adjusted to ensure sterility. The maximum allowable error was obtained from ISO8655 standards. Accuracy ranges for manual pipettes were obtained from various manufacturers of multi-channel pipettes.

Anticipating that there would be a need to vary the pipetting air-gap in the future to accomodate different operating volumes, we wished to develop a procedure that would enable quick and reliable determination of calibration parameters for the pipette tips. While automated gravimetric methods have been explored in the past for calibrating liquid handling pipettes, these would require the presence of a specialized, on-deck high-accuracy balance with minimal air-flow to prevent evaporation^1^, which may not be available on most liquid handling decks. We expected that the volume estimates calculated using the photometric standard could be used calibrate the pipettes. Upon analysis, we found a strong linear correlation between the pipetted volumes and the expected volumes within three different volume ranges - high (50 μL - 200 μL), mid (10 μL - 50 μL), and low (3 μL - 10 μL). Hence, we programmed the liquid handler to pipette eight different levels of the photometric standard within the three volume ranges in triplicate (Figure 4b). To enable automated processing of the data, we developed a python based script that accepts the absorbance data of the photometric standard along with the layout of the microplate used for calibration to determine the pipetting error for each volume pipetted. The script is then made to generate calibration parameters by performing a linear fit between the programmed/ expected volume and the actual pipetted volume. Using these parameters it is possible to determine the volume that needs to be programmed into the liquid handler for a required volume to be pipetted. Using these new calibration parameters, we analyzed the pipetting accuracy for each of the custom volume ranges with the increased air-gap. We found that our photometric calibration procedure reduced the deviation for all pipette tips significantly and brought them well below the maximum acceptable limits and within the ranges guaranteed by pipette manufacturers for multi-channel pipettes (Figure 4c). By using only on-deck components for calibration and a python script to automatically calculate calibration parameters, we were able to reduce the time required for calibrating each volume range to about 10 minutes. This protocol and the python script can be easily adapted to calibrate a wide variety of liquid handlers and conserve accuracy when changing the pipetting parameters.

### Maintaining sustained anaerobic environments in microplates

Having established protocols to eliminate contamination and calibrate pipettes, we aimed to investigate our platform’s ability to accelerate the ‘test’ phase of the DBTL cycle in metabolic engineering. As mentioned before, we were particularly interested in developing protocols for anaerobic phenotyping of enzymes and microbial strains in microplates due to the oxygen limiting nature of most high density fermentation processes. Short enzyme assays under anaerobic conditions can be achieved with relative ease through the addition of the oxygen scavenging enzymes such as glucose oxidase or Oxyrase along with suitable substrates^12^ in each well of the microplate. However, accurate phenotyping of microbial strains under anaerobic conditions using such enzymatic de-oxygenation would be challenging due to the need for glucose or other substrates for the enzymes to function. This would hinder accurate quantification of these metabolites after fermentation, resulting in incomplete carbon balances. Therefore, we decided to to use an anaerobic chamber to remove oxygen from the microplate by subjecting it through cycles of vacuum and flushing with nitrogen gas.

While anaerobic chambers are excellent for expelling oxygen from microplates, they require additional sophisticated equipment to control humidity. Without humidity control, the evaporation rates within anaerobic chambers are quite high, resulting in loss of media volume. Upon culturing different *E. coli* strains within the anaerobic chamber, we found that the rates of evaporation were so high that accurate measurements of cell density could not be made even though the duration of our fermentations were quite short (Supplementary Figure S3). As a possible solution, we examined the sealing efficacy of various adhesive films to sustain the anoxic conditions generated within the anaerobic chamber for fermentations outside. To measure of oxygen penetration into the microplate, we calculated biomass yields (ratio of final to initial biomass, measured as absorbance at 600 nm) of wild type *E. coli* (MG1655) grown to saturation in a rich defined medium within each well. Since *E. coli* grows faster under aerobic conditions, we should expect a consequent higher yield in wells that have increased oxygen penetration and low yields where anoxic conditions were sustained. As expected, in our control with a gas permeable film, we found a relatively high median biomass yield - characteristic of high oxygen penetration (Figure 5a). The use of a microplate lid with anaerobic adhesive tape did not offer much improvement in the seal, with only a modest decrease in the median biomass yield. The aluminium and polyester seals (typically used in PCRs) offered a significant improvement in the seal, with the polyster film being able to reduce the variability amongst wells as well. However, upon analysis of the biomass yield distribution within the microplates, we found clear patterns of enhanced growth in certain areas, likely resulting from improper sealing and heterogeneous oxygen concentrations(Figure 5a and Supplementary Figure S2). Hence, the use of a film would inevitably lead to heterogenity in cellular phenotypes in addition to increased throughput times due to the need for manual sealing of each microplate.

**Figure 5:**
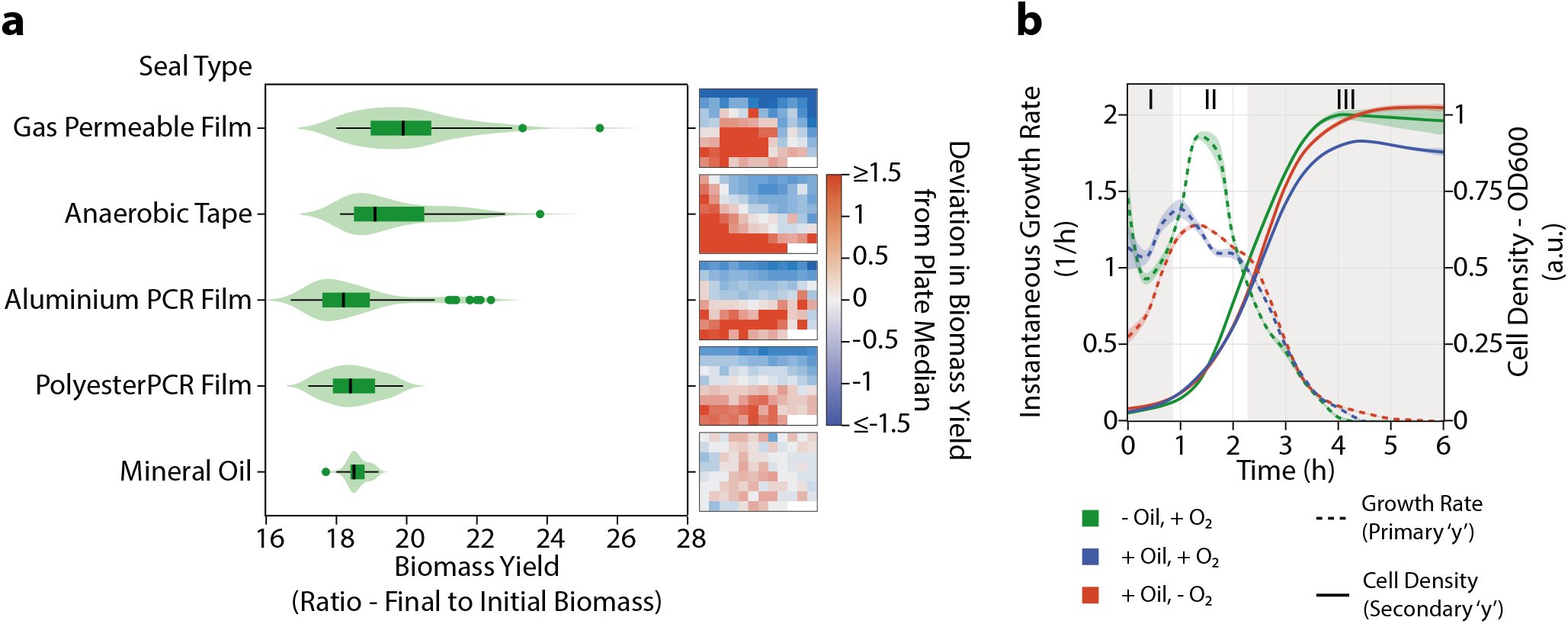
Establishing anaerobicity in 96-well microplates. **a.** Effectiveness of various seals in preventing oxygen penetration into microplates containing *E. coli* MG1655 in RDM, sealed within an anaerobic chamber. The biomass within each microplate are represented as violin plots. To the right of each violin plot, the distribution of biomass yields are represented as heatmaps showing deviation of the biomass yields from the median biomass yield within that plate. **b.** Time-course showing cell density and instantaneous growth rate of *E. coli* MG1655 in RDM with and without a layer of oil in the presence of oxygen and with a layer of mineral oil inside an anaerobic chamber.

Alternatively, a layer of mineral oil (50 μL), pipetted on top of the microbial culture in each well offered a homogeneous gas exchange profile, evidenced by the tight distribution of biomass yield (Figure 5a and Supplementary Figure S2). The mineral oil was also successful at completely eliminating loss of media during the fermentation within the anaerobic chamber, restoring the ability to monitor growth accurately (Supplementary Figure S3). In order to ensure that the growth profiles of *E. coli* are only affected by the resulting oxygen transfer and not directly by the mineral oil, we examined the growth of four different strains of *E. coli* with and without the layer of mineral oil, inside and outside the anaerobic chamber (Figure 5b and Supplementary Figure S3). We were able to clearly distinguish three different regimes in all the growth profiles - (I) an initial regime where dissolved oxygen in the media is used, indicated by the relatively higher growth rates of cells grown outside the anaerobic chamber, (II) an intermediate regime where the cells without the layer of mineral oil outside the anaerobic chamber are able to grow at accelerated rates due to increased oxygen transfer, and (III) a final growth phase where all the cells grow at similar rates due to no oxygen transfer due either to high cell densities or to the layer of mineral oil. It can be inferred from growth regimes (I) and (III) that the mineral oil does not directly impair or assist the growth of the strains but only controls the rate of gas exchange. Hence, it is suitable to maintain anoxic growth within an anaerobic chamber for extended durations with minimal loss of media due to evaporation.

### Case Study 1: Applying the liquid handling platform for an anaerobic enzymatic screen

As a preliminary validation of our high throughput phenotyping platform, we sought to perform an anaerobic activity screen of the enoate reductase enzyme YqjM from *Bacillus subtilis* (*Bs*-YqjM). This enzyme belongs to the family of old yellow enzymes (EC 1.6.99.1) which are broadly known as enoate reductases. They use non-covalently bound flavin mononucleotide (FMN) to catalyze the reduction of double bonds found in *α,β*-unsaturated aldehydes and ketones using NADPH or NADH as electron donors^13^. The ability of *Bs*-YqjM and other enoate reductases to reduce -ene groups is important for the catalysis of chemical commodities such as muconic acid to adipic acid (a pre-cursor to nylon). However, the activity of *Bs*-YqjM enzymatic activity is known to be supressed in the presence of oxygen under aerobic conditions due to a prominent background reaction where electrons from NADPH are transferred to dissolved molecular oxygen in the buffer. In contrast, its activity is markedly increased under anaerobic conditions where electrons are instead donated to its target -ene substrates^40^. For the 2-cyclohexen-1-one substrate, Bs-YqjM was reported to have a K*_M_* value of 0.3-0.6 mM under anaerobic conditions created using a glucose-glucose oxidase system, which consumes the dissolved molecular oxygen in the buffering solution to simulate completely anaerobic conditions.

To demonstrate the use of an automated LiHa platform for performing anaerobic assays, we purified BsYqjM and assayed its activity for 2-cyclohexen-1-one by monitoring changes in the absorbance at 340 nm due to NADPH oxidation. After calibration of the tips for smaller volumes in the 3-10 μL range, we observed a K*_M_* value of 0.35 ± 0.06 mM using the automated platform (Figure 6a). In comparison, we performed the same assay manually and observed a K*_M_* value of 0.33 ± 0.4 mM. The similarity of these K*_M_* values to each other and to published literature values suggested that the LiHa platform could be used to automate the preparation of screens, such as those to determine the optimal pH for maximum activity. Towards this end, we determined Bs-YqjM’s activity across pH 2.2 – 8 using the liquid handler (Figure 6b). We found that BsYqjM operates optimally at pH 5-6, which aligns with previously reported results that Bs-YqjM prefers slightly acidic conditions^40^.

**Figure 6:**
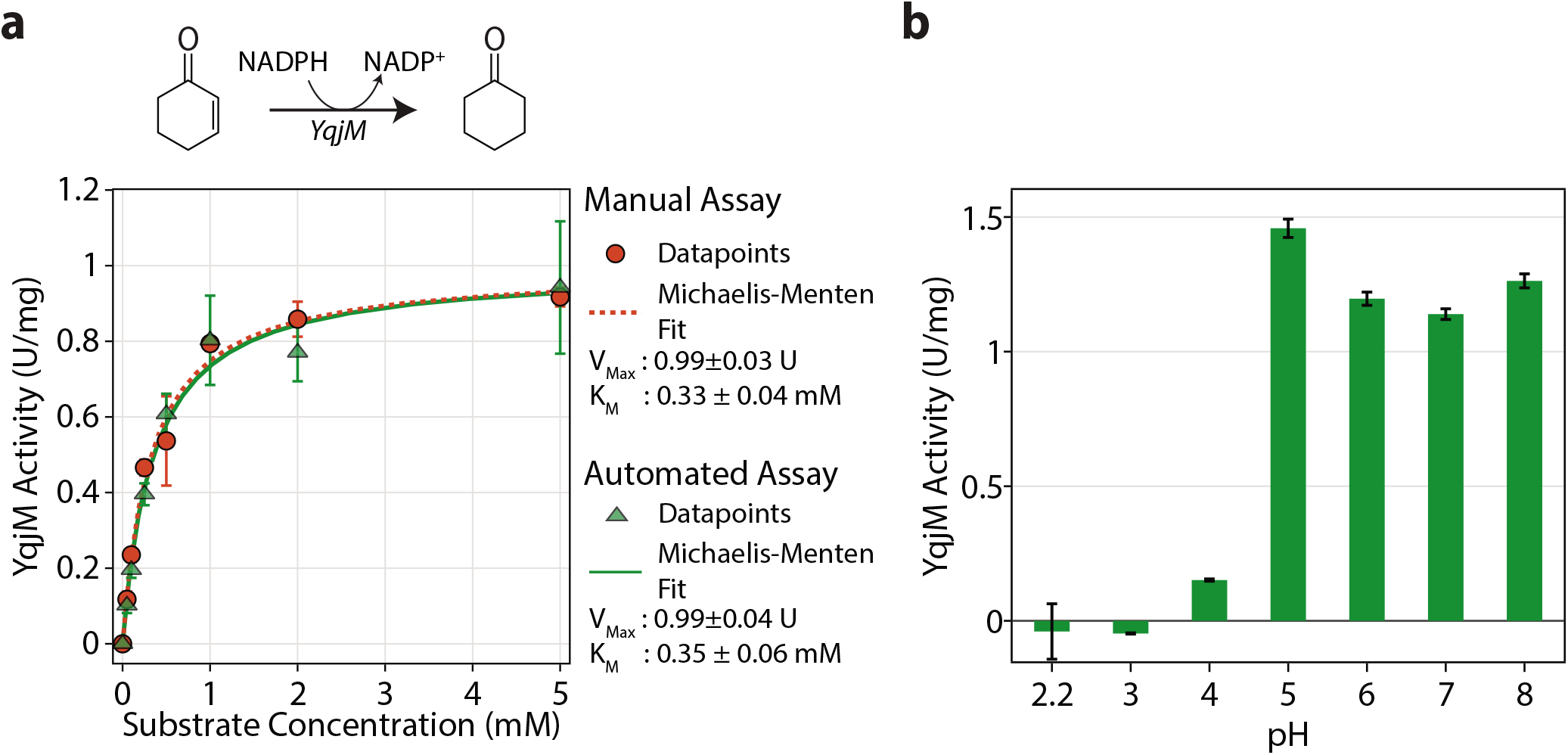
Anaerobic enzymatic screen. **a.** Enzymatic activity of YqjM on 2-cyclohexen-1-one determined manually and by the liquid handler. Enzyme activity is represented in units of μmol/min. **b.** Effect of pH of the medium on the activity of YqjM on 2-cyclohexen-1-one.

### Case Study 2: Scaling down anaerobic microbial phenotypes from pH controlled bioreactors to microplates

Having assessed the efficacy of our system in determining enzyme kinetic parameters under anaerobic conditions, we wished to investigate the applicability of a fixed-tip liquid handling system for a high-throughput characterization of microbial phenotypes under anaerobic conditions. While it is possible to rapidly cultivate microbial strains using our platform, the possible deviation of phenotypes at increasingly larger scales is a cause for concern, resulting in ambiguity of the strains to be chosen for further screening. Previous studies examining scaling considerations have primarily investigated the difficulty of improving oxygen transfer rates within the wells of microplates^25,54^. However, since we are interested only in anaerobic environments, oxygen transfer rates may not play a key role in determining phenotypes. Rather, the concentration of substrate, pH, and other media conditions could be the determining factors. Hence, as a second test case to validate our platform, we investigated the ability to scale-down microbial phenotypes observed in pH controlled 500 mL bioreactors to 96 well microplates under anaerobic conditions. To this end, we examined the growth and metabolite profiles of four strains of *E. coli* - MG1655 and its lactate overproducing deletion mutant, MG1655 Δ(*adhE, pta*) at three different stages of adaptive laboratory evolution (denoted Δ(*adhE, pta*)-D1, D28 and D59 to represent the duration of adaptive laboratory evolution in days) ^14^. These strains were chosen because of the expected difference in their anaerobic phenotypes. During anaerobic growth, *E. coli* undergoes mixed acid fermentation due to the non-availability of oxygen as a terminal electron acceptor to produce ATP and regenerate the redox cofactors NAD and NADP. Instead, *E. coli* produces a mixture of formate, acetate, ethanol, lactate, and small quantities of other organic acids as terminal fermentation products (Figure 7a), with acetate, ethanol, and formate being preferred products due to higher energy yields. Due to deletions around key fermentation reactions involved in acetate and ethanol production (pta and adhE respectively), the deletion mutants used in our study are expected to show high lactate yields. Further, because these strains are products of adaptive laboratory evolution, those strains at a later stage of evolution are expected to show increased growth rates.

**Figure 7:**
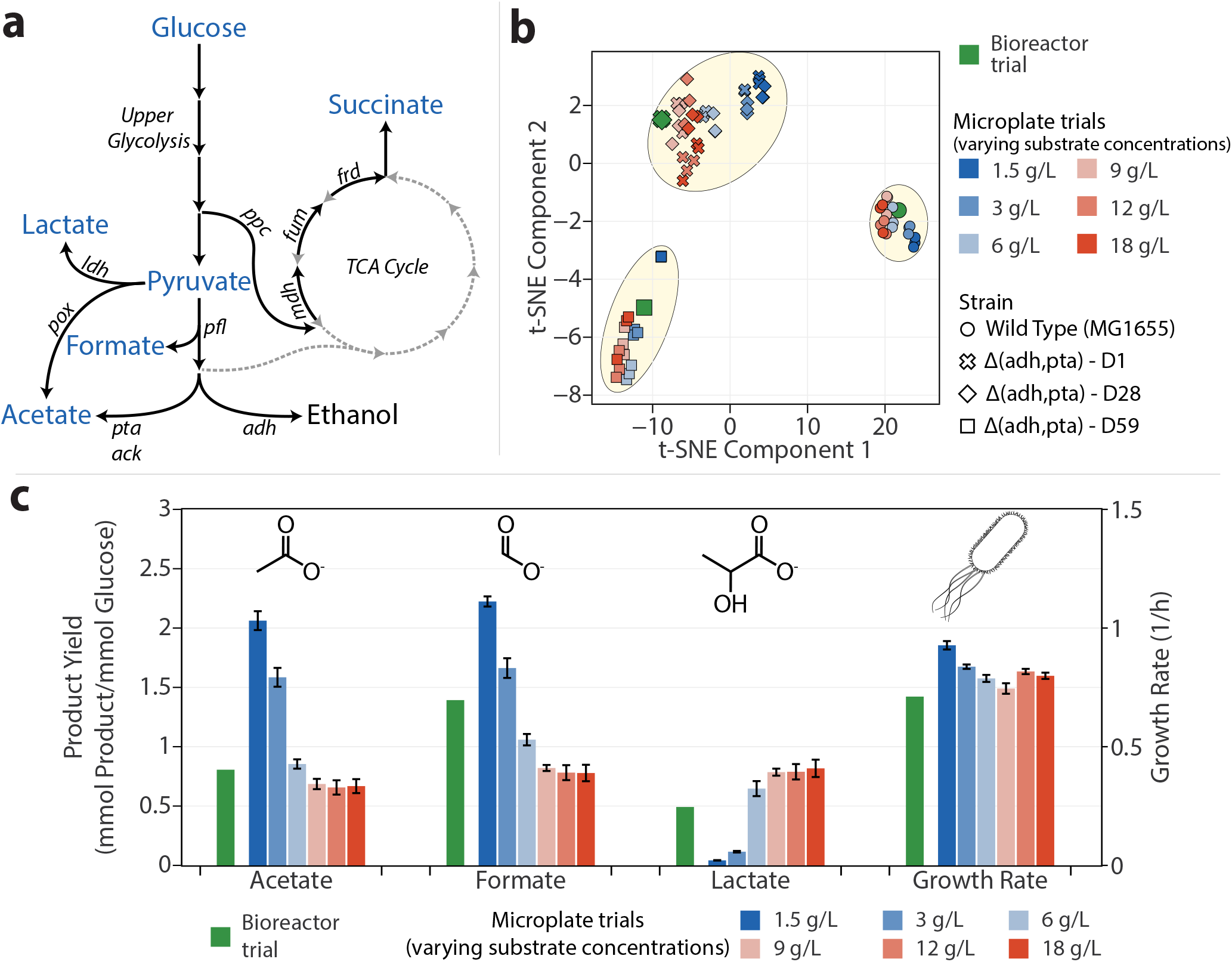
Comparison of *E. coli*’s anaerobic phenotype in bioreactors and microplates. **a.** Schematic showing typical fermentation pathways in *E. coli*. Typical products of mixed acid fermentation on glucose are shown in the pathway along with key fermentation reactions shown in italics. The metabolites measured in this study are shown in blue. **b.** Microbial phenotypes reduced to two components through t-distributed stochastic neighbors embedding (t-SNE) performed on the metabolite (acetate, formate, lactate, pyruvate, and succinate) yields and growth rates of *E. coli* strains grown in rich defined media in a bioreactor and microplates with different initial concentrations of substrate (glucose). Cluster boundaries were drawn manually for illustrative purposes. **c.** A comparison of WildType *E. coli*’s growth rate and metabolite yields on glucose obtained from a bench-top 0.5 L bioreactor and 96-well microplates with different initial concentrations of substrate (glucose).

To compare the metabolic state of the different strains grown in a bioreactor and microplates, we calculated the growth rates and yields of five different products of fermentation on glucose towards the end of the exponential phase of growth(Supplementary Figure S7). The deletion mutants grown in microplates showed good agreement with the bioreactor phenotype as is, possibly due to the elimination of the most prominent fermentation modes - acetate and ethanol production. However, the wild type strain showed pronounced phenotypic differences in the microplate, producing significantly lower levels of formate. It appeared that more carbon flux was directed towards lactate production than formate production in the microplates, resulting in less energy efficient fermentation and therefore, reduced growth rates. In order to eliminate the possibility of residual dissolved oxygen in the media causing aberrant phenotypes and lower formate yields, we examined the effect of adding the reducing agents - 1 mM cysteine, 1 mM dithiothreitol (DTT), and 8 mM sodium sulfide to scavenge any residual oxygen and maintain reducing conditions within the media (Figure 7b and Supplementary Figure S7). Higher concentrations of sodium sulfide were chosen because previous experiments at the 1mM level showed no visible differences in the phenotype. To better visualize and compare the overall phenotypic differences resulting from the different strains and media conditions, we performed a dimensionality reduction of the seven analytes (growth rate and yields of acetate, formate, lactate, pyruvate, succinate and biomass on glucose) through principal component analysis (PCA) (Supplementary Figure S6). Upon analysis of the scores of each experimental trial on the first two principal components, the bioreactor trial for the wild-type strain resulted in phenotypes which could not be replicated in microplates since the bioreactor trials seemed to be isolated from the clusters formed by the microplate trials. Further, PCA indicated that residual oxygen may not an issue since the addition of reducing agents did little to alter the phenotypes. Examining the individual analytes (Figure S7), we found that the addition of cysteine at 1 mM did not alter the metabolite and growth profiles significantly for any of the strains. The addition of DTT showed a decrease in the yield of nearly all products including biomass for all strains, indicating that it could be inhibitory to the cells. Interestingly, the addition of sodium sulfide seemed to push the metabolic state slightly towards that observed in the bioreactor, with increased growth rates and acetate yields but lower lactate yields. However, since we did not observe similar phenotypes using the other reducing agents, we hypothesized that this difference could be due to the basic nature of sodium sulfide, which would result in longer fermentation times and therefore a different metabolic profile. We confirmed this by growing *E. coli* at a higher starting pH, resulting in longer fermentation duration, and similar trends in the metabolite yields and growth rates as observed in the addition of sodium sulfide.

Hence, we concluded that our platform resulted in complete anaerobicity of the media and it was not dissolved oxygen that was affecting the metabolic state of the cells. It appeared that the pH and consequently, the fermentation duration played a more important role in determining the phenotype of the wild-type strain, as expected. The implementation of pH control in microplates requires specialized microplates with base delivery systems or mini-bioreactors, which would greatly increase operational costs^15,50^. We proposed that varying initial glucose concentrations would offer a crude yet inexpensive means to alter the duration of fermentation, thereby limiting pH change, and consequently, impact the phenotypes of all strains. Therefore, we grew the *E. coli* strains with different starting concentrations of glucose to examine this effect and determine glucose concentrations that allowed the phenotype of the wild-type strain observed in the bioreactor trial to be replicated in microplates (Figure 7c and Supplementary Figure S9). At high initial glucose concentrations, all strains showed increased lactate yields and reduced biomass, formate and acetate yields on glucose. Specifically, for the wild type strain, this indicates that a significant portion of the carbon flux is directed towards lactate production with reduced flux through *pfl, pta*, and *adh*, resulting in less efficient fermentation and reduced growth rates. However, at lower substrate concentrations, the overall fermentation duration and consequently, the pH change during the fermentation decrease. This results in less overflow of carbon flux towards lactate and increased yields of biomass, acetate and formate, with almost no lactate and maximal formate, acetate and growth rates at the lowest concentrations analyzed. Performing the same dimensionality reduction through PCA as described previously, we found that varying initial glucose concentrations significantly alters the overall phenotypes exhibited by the cells, as shown by the spread of the scores of each experimental trial in the principal component space (Supplementary Figure S8). Interestingly, several microplate trials with overall phenotypes very close to their bioreactor counterparts for each strain were observed. Particularly, the wild-type strain seemed to be closest to the microplate trial starting with 6 g/L of glucose. The other strains seemed less impacted by high initial glucose concentrations and showed good agreement with the bioreactor phenotype even at high glucose concentrations.

While these results indicate that phenotypes observed in bioreactors can be reasonably replicated in microplates by varying initial substrate concentrations, the exact value for each strain may not be the same, as seen here. Further, the optimal glucose concentration for each strain cannot be determined a priori, which may lead to ambiguity in determining better performing strains to be chosen for scale up. Hence, we wished to investigate dimensionality reduction techniques, using which strains showing similarities at the bioreactor scale could be clustered together while segregating those that showed significant differences. Our dataset from the experiments varying initial glucose concentrations was ideal for this purpose since we observed an array of different phenotypes at the microplate scale for the same strain. Further, the mutant strains - Δ(adh, pta)-D1 and Δ(adh, pta)-D28 showed very similar phenotypes at the bioreactor scale. As seen previously (Supplementary Figure S8), principal component analysis was only partially successful in this effort - while most trials with the D1 and D28 strain exp appeared in the same cluster, trials with the D59 strain also occurred very close to them. Moreover, the wild-type strains could not form a single cluster, possibly due to the large variability in the metabolite yields. Hence, a two-dimensional PCA alone cannot be used to determine strains that would show similar performance at larger scales, possibly due to omitting the variance expained by the other principal components. A relatively new dimensionality reduction algorithm - t-distributed stochastic neighbors embedding, which recreates the probability distribution of entities in a higher dimensional space to two dimensions, has been found to be successful at clustering similar entities when a large number of dimensions are involved^49^. Particularly, it has found use in analyzing single cell transcriptomic data. Eventhough our dataset is comprised of only 6 dimensions i.e. the yields of five metabolites and the growth rates, we proposed that tSNE could potentially be successful at clustering similar performing strains in a reduced dimensional space, particularly due to its use of non-linear dimensionality reduction. Remarkably, a tSNE model fit to our glucose varying data showed near perfect clustering of strains showing similar performance at the bioreactor scale (Figure 7b). Specifically, all microplate trials from the wild-type strains and the Δ(adh, pta)-D59 strain were resolved into their individual clusters in spite of the visible differences in the phenotypes of individual trials. The two mutants Δ(adh, pta)-D1 and Δ(adh, pta)-D28 that showed similar performance at the bioreactor scale were resolved into a single cluster. These results indicate that tSNE could be used effectively to shortlist strains for analysis at larger scales, since it is able to effectively segregate strains showing markedly different phenotypes. Therefore, while initial glucose concentrations affect the phenotypes of microbial strains at the microplate scale significantly, the use of dimensionality reduction techniques such as tSNE could be used to resolve these differences and identify overall phenotypic differences between strains.

## Conclusions

We have seen that our automated platform is able to rapidly and effectively set up microplate experiments to phenotype enzymes and microbial strains. The automation of such routine metabolic engineering workflows greatly expands the number of different strains/enzymes and media conditions that can be examined, resulting in large experimental datasets that can assist strain design. With machine learning applications in metabolic engineering becoming more prevalent, there is an urgent need to develop tools and protocols for accurate and reproducible phenotyping strains and enzymes at smaller scales. Automated systems are uniquely suited for this task since they eliminate human error and require standardized protocols to function. Furthermore, recent efforts toward developing robot programming languages that allow for the development of cross-platform protocols enable relatively easy implementation of complex laboratory workflows^33,34,55^.

While automation can enhance experimental throughput, conducting experiments at accelerated rates also increases operational costs and the amount of laboratory waste generated due to the number of pipette tips and other labware used. Laboratory plastic waste has become a major concern in the current era of high-throughput experimentation^3,30,48^. It is quite ironic that the same research labs that work on developing microbes for sustainable production of chemicals end up generating several million tonnes of plastic waste in the process. Through the development of effective and fast decontamination protocols, we eliminated the need for plastic pipette tips while maintaining experimental throughput. Disregarding repeated and failed experiments, we estimate that nearly 4000 pipette tips would be required to complete the two case studies examined in this work if they were done manually or using a disposable tip liquid handling platform. Further, the automated pipette calibration protocol developed here enables the quick setup of a broad range of liquid handling systems for different pipetting programs and would also assist in routine maintenance without the need for additional expensive equipment.

One concern with phenotyping microbial strains in microplates is the inability to replicate the mixing regimes, oxygen transfer and other physical characteristics of fermentation observed in larger pH controlled bioreactors. These considerations are better addressed in miniature bioreactors that have been designed to be small scale replicas of bench-top bioreactors. Nevertheless, by leveraging the enhanced throughput of microplate experiments, we were able to analyze the effect of a large number of media conditions on the cellular phenotypes in a relatively short period of time. Consequently, we were able to identify glucose concentrations that restricted fermentation durations and thereby, reasonably reproduce bioreactor phenotypes in microtiter plates under anaerobic conditions. Furthermore, modern dimensionality reduction and data visulalization techniques such as tSNE could be used in conjunction with microplate experiments as scale-down models to assist in choosing strains for scale-up. We believe that since microplates offer higher experimental throughput at very low costs, our platform will serve as an effective and representative screen before moving on to larger scales. Furthermore, integration of our data analysis pipeline - IMPACT with the strain testing pipeline has enabled the visualization and analysis of large datasets that emerge as a consequence of our platform, and will accelerate future strain design endeavours. While successful at anaerobic phenotyping, we believe that the experimental protocols described in this study are broadly applicable to various liquid handling platforms for a wide range of applications and this work will assist the development of sustainable automated high throughput experimental platforms.

## Materials & Methods

### Enzymes, Strains and Experimental Medium

Wild type *Escherichia coli* strain K-12 MG1655 was used to detect contamination during the development of our decontamination protocol. The wild type *Escherichia coli* strain K-12 MG1655 and its mutants harboring deletions of the genes *adhE* and *pta* at three different stages of adaptive laboratory evolution^14^ (denoted Δ(adhE, pta)-D1, Δ(adhE, pta)-D28, and Δ(adhE, pta)-D59 to reflect duration of adaptive laboratory evolution in days) were used to examine the efficacy of our phenotyping platform. The enoate reductase enzyme yqjM (UniProt: P54550) from Bacillus subtilis strain 168 was used for the anaerobic screen.

Lysogeny Broth (LB) media was used to prepare bacterial starter cultures in all cases. Strain phenotyping experiments were conducted in a rich defined medium (RDM) composed of a carbon source (D-glucose at various concentrations), salts (3.5 g/L KH_2_PO_4_, 5 g/L K_2_HPO_4_, 3.5 g/L (NH_4_)_2_HPO_4_, 1 mM MgSO_4_, 0.1mM CaCl_2_), 1 mM 3-morpholinopropane-1-sulfonic acid (MOPS), amino acid supplements (0.8 mM alanine, 5.2 mM arginine, 0.4 mM aspargine, 0.4 mM aspartate, 0.1 mM cysteine, 0.6 mM glutamate, 0.6 mM glutamine, 0.8 mM glycine, 0.2 mM histidine, 0.4 mM isoleucine, 0.8 mM leucine, 0.4 mM lysine, 0.2 mM methionine, 0.4 mM phenylalanine, 0.4 mM proline, 10 mM serine, 0.4 mM threonine, 0.1 mM tryptophan, 0.2 mM tyrosine, and 0.6 mM valine), nucleotide supplements (0.1 mM each of adenine, cytosine, guanine, and uracil), and vitamin supplements (0.01 mM each of thiamine, calcium pantothenate, *p*-aminobenzoic acid, *p*-hydroxybenzoic acid, and 2,3-dihydroxybenzoic acid) - adapted from the defined media composition described previously^36^. All media components were sterilized either by autoclaving or filter sterilization. Stocks of cysteine, dithiothreitol (DTT), and sodium sulfide for use as reducing agents to maintain anaerobicity in the media were prepared at a concentration of 0.2 M. The stocks were sparged gaseous nitrogen through the solutions for 15 minutes to eliminate dissolved oxygen, followed by sterilization.

12% sodium hypochlorite (Bioshop SYP001.1) and 95% ethanol were diluted to required concentrations to prepare disinfectants for the decontamination protocol. Aqueous solutions of potassium dichromate (0.4 mM, 1mM, and 2mM) were prepared to detect pipetting accuracy and calibrate the liquid handling system. The polyurethane gas permeable film (Diversified Biotech BEM-1), polyester PCR film (Bio-Rad MSB1001), and aluminized foil (Bio-Rad MSF1001) were used to seal 96 well microplates (Corning 353072) containing *E. coli* cultures to investigate anaerobicity. Mineral oil (BioShop MIN444) was used to prevent evaporation in anaerobic chambers where required.

### High throughput phenotyping platform

The phenotyping platform described in this study was comprised of a Tecan Freedom Evo 100 base fitted with a Tecan fixed-tip liquid LiHa (liquid handling) arm, a Tecan RoMa (robotic manipulator) arm, a QInstruments Bioshake 3000-T microplate heater-shaker, an Agilent microplate centrifuge, a Tecan Infinite M200 plate reader, and a Tecan Te-VacS vacuum filtration module. Communication with the various modules and all automation scripts were set up on Tecan’s EvoWare 2.7 platform.

### Enzyme Purification for Anaerobic Screen

The gene encoding yqjM was cloned under the T7 promoter in-frame with an N-terminal 6x HisTag of the p15TvL expression vector (AddGene: 26093) using the In-Fusion@HD EcoDry kit, and then transformed into LOBSTR BL21(DE3) Escherichia coli. Starter cultures for yqjM were grown from glycerol stock in lysogeny broth (LB) media with ampicillin (100 μg/mL) for 16 hrs at 37 °C with shaking. Then, expression cultures were started in 1L Terrific Broth media with ampicillin (100 μg/mL) and a 1% v/v inoculant of the starter culture, followed by growth for 4 hrs at 37 °C and induction with 0.4 mM IPTG. The expression culture was then transferred to 16 °C and grown for 16 hrs with shaking, pelleted with centrifugation, and transferred to vials for one freeze-thaw cycle at −20 °C. Frozen cell pellets were thawed and resuspended in binding buffer (10 mM HEPES, 500 mM NaCl, 5 mM imidazole, pH 7.2) to a final volume of 50 mL, followed by addition of 0.25 g lysozyme. Cell pellet mixtures were sonicated for 25 min and clarified by centrifugation. The supernatant was applied to a cobalt-charged NTA resin pre-equilibrated with binding buffer in a gravity-column set-up. Bound proteins were cleansed with 120 mL of wash buffer (10 mM HEPES, 500 mM NaCl, 25 mM imidazole, pH 7.2) and collected with 4 mL elution buffer (10 mM HEPES, 500 mM NaCl, 250 mM imidazole, pH 7.2). Protein concentrations were determined by Bradford assay to be 4.3 mg/mL (120 μM), and protein purity was determined by SDS-PAGE analysis and densitometry to be 99%. A molar equivalent of flavin mononucelotide (FMN) was loaded into YqjM prior to transfer into a 10 kDa MWCO dialysis bag for dialysis in 1 L dialysis buffer (40 mM HEPES, pH 7.5) at 4 °C with gentle stirring for 24 hrs. YqjM was then flash frozen drop-wise in liquid nitrogen before storage at −80°C.

### NADPH Assay for Determination of Anaerobic YqjM Activity

The glucose oxidase type VII-S from *Aspergillus niger* was used to remove oxygen from enzyme screen reactions using D-glucose as the substrate. Working concentrations of 2-cyclohexen-1-one (substrate), β-NADPH tetrasodium salt (indicator), glucose oxidase type VII-S from Aspergillus niger, and glucose were prepared in 40 mM HEPES at a pH of 7.5 to assay yqjM activity. Assays were set up in a 96 well microplate using the liquid handler and consisted of 0.15 mM NADPH, 10 u/mL glucose oxidase, 20 mM glucose, and 15 nM YqjM. The substrate, 2-cyclohexen-1-one was then added at required concentrations along with the activity buffer to make each assay up to a volume of 200 μL. The pH gradients were prepared using McIlvaine buffers with appropriate ratios of 0.2 M Na_2_HPO_4_ and 0.1 M citric acid which replaced the activity buffer. Salt gradients were prepared by adding appropriate concentrations of NaCl and KCl to the activity buffer.

YqjM activity was determined by measuring NADPH concentrations in triplicate using kinetic reads performed using a Molecular Devices SpectraMax M2 spectrophotometer at 35°C at an absorbance wavelength of 340 nm with shaking before and in between kinetic reads. The volumetric activities (μmol min-1 mg-1) were calculated using NADPH’s extinction coefficient of 6.3 mM^−1^ cm^−1^ and a height of 0.56 cm. The obtained activity data was fit to a Michaelis-Menten curve to obtain *K_M_* and *V_Max_* through non-linear regression using optimization tools in the python package - scipy^53^.

### Determination of microbial phenotypes in microplates

*E. coli* strains streaked on LB-agar plates were used to prepare starter cultures for the scaled down phenotyping experiments. The strains were inoculated in LB media supplemented with 1% glucose in 96 well microplates and grown overnight at 37°C with constant shaking at 250 rpm. Glucose was added to the starter cultures to eliminate the need for an intermediate adaptation culture in the experimental media - RDM (Supplementary Figure S4). The microplates containing the overnight precultures were then transferred to the liquid handling platform for processing. All following steps were automated on the liquid handling platform.

First, to remove traces of fermentation products and spent media from the strains, the pre-cultures were harvested by centrifugation at 3000 g for 10 minutes and washed with RDM lacking carbon source 2 times before being resuspended in the experimental RDM medium consisting of the carbon source and any required supplements. Then, the cell density of each well was determined by measuring the absorbance at a wavelength at 600 nm on a Tecan Infinite M200 plate reader and cells were then diluted to a cell density of 0.05 with appropriate media to a final volume of 150 μL to normalize all wells to the same starting OD.

After this, the plate was removed from the liquid handling platform, taken through cycles of vacuum and flushing with nitrogen gas, and transferred into an anaerobic chamber filled with N_2_ gas. The cultures were then covered with a 50μL layer of laboratory grade mineral oil (BioShop MIN444) to prevent evaporation. To ensure anaerobic conditions throughout the fermentation, the cells were grown within the anaerobic chamber at 37°C and constant shaking in a Molecular Devices SpectraMax M2 platereader which also recorded the cell density periodically.

After the cells finished growing (about 8h), the microplate was removed from the anaerobic chamber and transferred to the liquid handler for HPLC sample preparation. The liquid handling platform was programmed to pipette the samples onto a 0.2 μm filter plate (Millipore MSGVN2210) for filtration. Samples were filtered at 400 psi for 60 s into a sample collection plate. Fermentation products were separated by passing the samples through an Aminex HPX-87H cation exchange column (BioRad 1250140) at a flow rate of 0.6 mL/min with 5mM H_2_SO_4_ as the mobile phase and 60°C column temperature. Metabolite concentrations were determined by monitoring the refractive index and UV absorbance (at 210 nm, 254 nm) of the eluent. The chromatograms were integrated using Chromeleon v7.

### Determination of microbial phenotypes in pH controlled bioreactors

*E. coli* strains streaked on LB-agar plates were used to prepare starter cultures by inoculation into 5 mL LB + 1% glucose. Cultures were then transferred to 50 mL sealed Falcon tubes for oxygen limitation. After overnight growth, cells were washed 3 times with RDM lacking carbon source before being transferred to 500 mL bioreactors (Applikon Mini) with 300mL of RDM with a glucose concentration of 2%. The media in the bioreactors was maintained anaerobic by sparging with nitrogen gas. pH was maintained at 7 within the bioreactor by continuous control using 10 M NaOH and the temperature was maintained at 37 °C. Samples for cell density and metabolite concentration measurements were withdrawn from the bioreactor periodically. Cell density was determined by measuring absorbance at 600 nm on a spectrophotometer (Thermo Scientific GENESYS20). Metabolite concentrations were determined through HPLC as described in the previous section after filtering the samples through 0.2 μm nylon filters.

### Data Analysis

Data analyses for all sections were conducted using Python on Jupyter notebooks. The jupyter notebooks used to generate figures and process data in this work are available on Github^42^. The python based data analysis library - pandas and plotting library - plotly were used extensively for all data analysis and visualization pipelines in this work^21,38^.

Microbial phenotypic data and growth curves were analyzed using the IMPACT Framework^51^. For the microbial phenotyping experiments, since time-course metabolite concentrations could not be obtained for the microplate trials, end-point metabolite concentrations were used to calculate yields. Hence, for a fair comparison with the microplate trials, yields for the bioreactor trials were calculated from metabolite concentrations obtained near the end of the exponential phase of growth. Growth rates for both bioreactor and microplate trials were determined from only the exponential phase of growth and were calculated as the specific biomass productivity (i.e. 1/[X]*d[X]/dt where [X] is the biomass concentration) and averaged over the required time-period. The sci-kit learn library was used perform principal component analysis (PCA) to reduce the dimensionality of scaled phenotype data (growth rates and yields of acetate, formate, lactate, pyruvate, and succinate on glucose) and enable easier phenotypic comparisons^39^. A number of components that explained at least 90% of the variance in the phenotypic data was chosen for PCA. Phenotypic data was scaled to unit variance and zero mean prior to PCA. Similarly, t-distributed stochastic neighbours embedding was also implemented from the sci-kit learn library. A perplexity that resulted in the most robust embedding was determined after iterating through several values. The learning rate (*ϵ*) that minimized the Kullback–Leibler divergence of the input data distribution and the resulting distribution was used. Regardless, other values of perplexity and learning rate resulted in similar results when an optimal solution was achieved.

## Supporting Information

Supplementary Table S1 in the Supporting Information contains details of reaction abbreviations used in Figure 7a. Supplementary Figures S1-S9 of Supporting Information are figures that have been referenced in the manuscript

## Author Contributions

KR helped formulate the study, developed automation protocols, performed experiments, developed the codebase, and wrote and edited the manuscript. NV helped formulate the study, developed automation protocols, and reviewed the manuscript. PD performed enzyme kinetic screens and wrote the manuscript. SG performed experiments. AY supervised the work and reviewed the manuscript. RM helped formulate teh study, supervised the work and reviewed the manuscript.

## Acknowledgements

The authors thank Prof. Stephen Fong (Virginia Commonwealth University, USA) for providing the lactic acid overproducing mutants used in this study. We also thank Prof. Po-Hsiang Wang (National Central University, Taiwan) for insightful discussions about maintaining anaerobicity in cell cultures. This work was financially supported by grants from Genome Canada, The Ontario Ministry for Research, Innovation, and Science, and the National Sciences and Engineering Research Council of Canada. KR would like to acknowledge funding from an Ontario Trillium Scholarship and a Mitacs Globalink Fellowship.

## Notes

### Competing Interest Statement

The authors have declared no competing interest.

