## Supplementary Table and Figures for "Automation Assisted Anaerobic Phenotyping For Metabolic Engineering"

**Table S1:** Names and formulae of reaction abbreviations mentioned in Figure 7 (main text). Metabolites in the reaction formulae are represented by their BiGG ID

| Abbreviation | Reaction Name | Reaction Formula |
|---|---|---|
| ack | Acetate kinase | ac_c + atp_c <=>actp_c + adp_c |
| adh | Alcohol/aldehyde dehydrogenase | acald_c + coa_c + nad_c <=>accoa_c + h_c + nadh_c |
| | | etoh_c + nad_c <=>acald_c + h_c + nadh_c |
| frd | Fumarate reductase | fum_c + mql8_c ->mqn8_c + succ_c |
| fum | Fumarase | fum_c + h2o_c <=>mal__L_c |
| ldh | Lactate dehydrogenase | lac__D_c + nad_c <=>h_c + nadh_c + pyr_c |
| mdh | Malate dehydrogenase | mal__L_c + nad_c <=>h_c + nadh_c + oaa_c |
| pfl | Pyruvate formate lyase | coa_c + pyr_c ->accoa_c + for_c |
| pox | Pyruvate oxidase | h2o_c + pyr_c + q8_c ->ac_c + co2_c + q8h2_c |
| ppc | Phosphoenolpyruvate carboxylase | co2_c + h2o_c + pep_c <=>h_c + oaa_c + pi_c |
| pta | Phosphotransacetylase | accoa_c + pi_c <=>actp_c + coa_c |

### Supplementary Figures

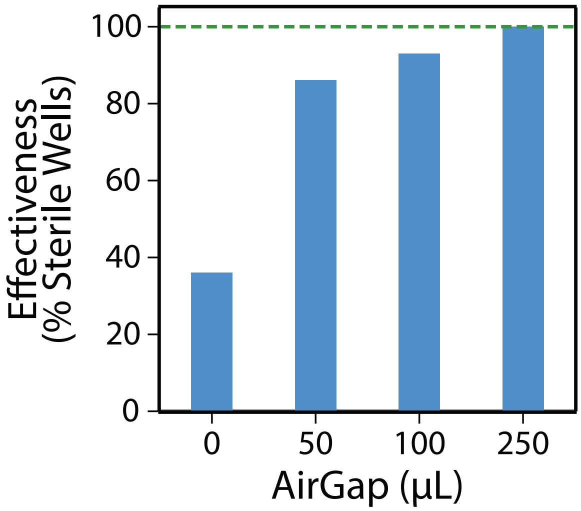


**Figure S1:** Change in sterility with air-gap (data consolidated from main text Figure 2d).

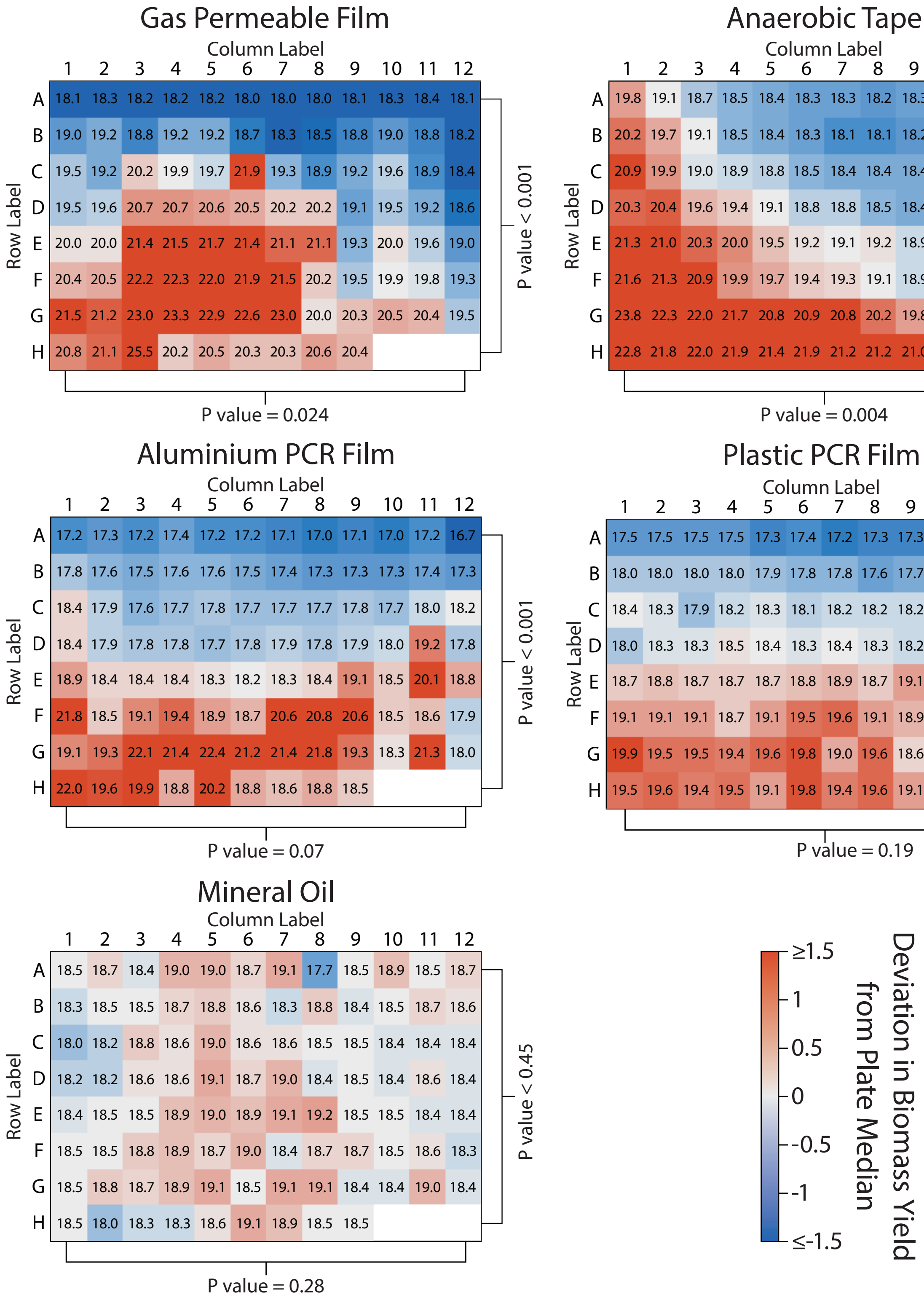


**Figure S2:** Distribution of biomass yields (ratio of final to initial biomass)of wild type *E. coli* MG1655 grown in Rich Defined Media with different seals. Yield values recorded in each well of the plates are shown, with a heatmap illustrating the deviation of the yield from the median value of the plate.

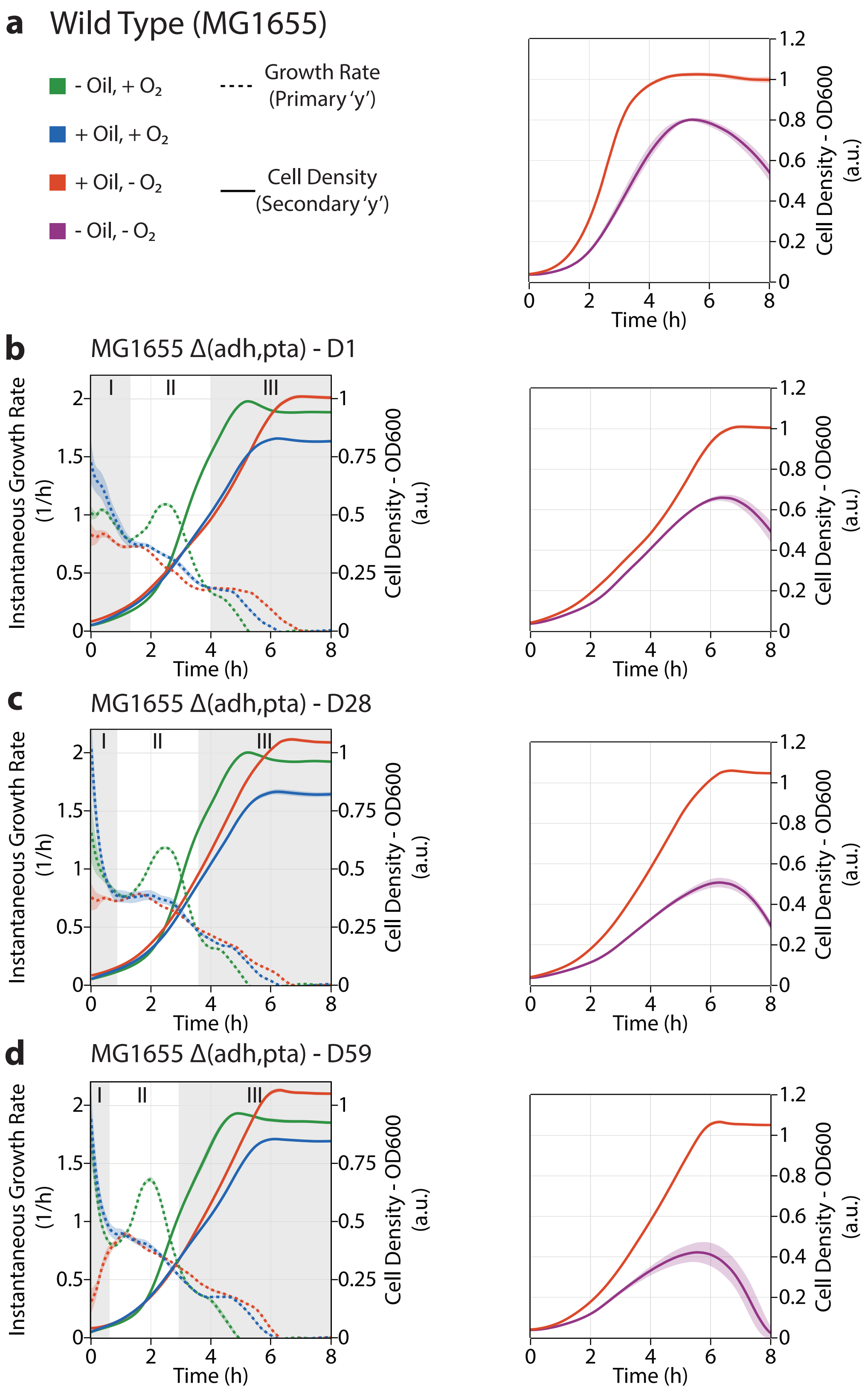


**Figure S3:** Time-course showing cell density and instantaneous growth rate of different *E. coli* strains (described in Materials & Methods) in RDM with and without a layer of oil in the presence of oxygen and with a layer of mineral oil inside an anaerobic chamber. Decreaase in absorbance of strains grown anaerobically without the oil is due to evaporation of culture media.

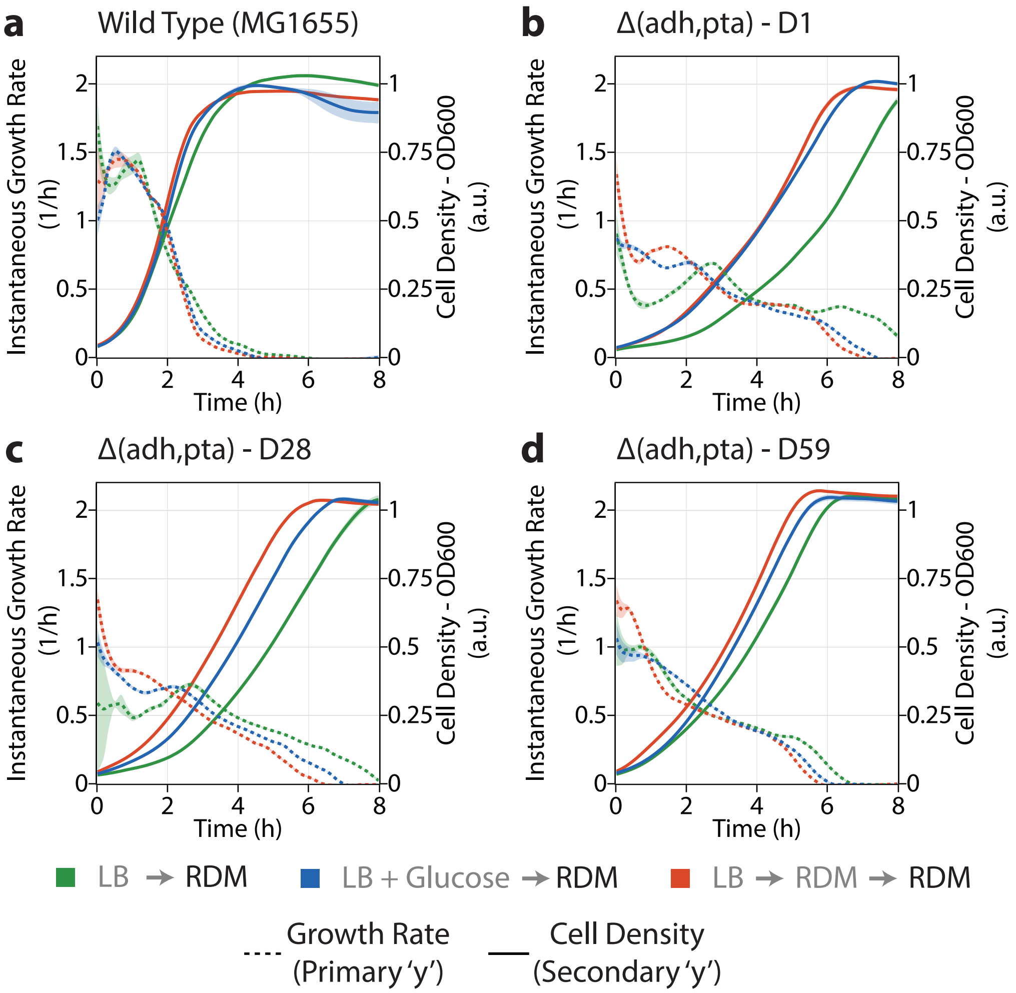


**Figure S4:** Time-course showing cell density and instantaneous growth rate of *E. coli* strains (described in Materials & Methods) with different pre-culturing strategies. For all strains, cells transferred from LB+glucose to RDM showed similar growth profiles to those with an intermediate adaptation transfer to RDM. In contrast, cells transferred from LB to RDM directly showed a longer lag phase and slower growth in all cases.

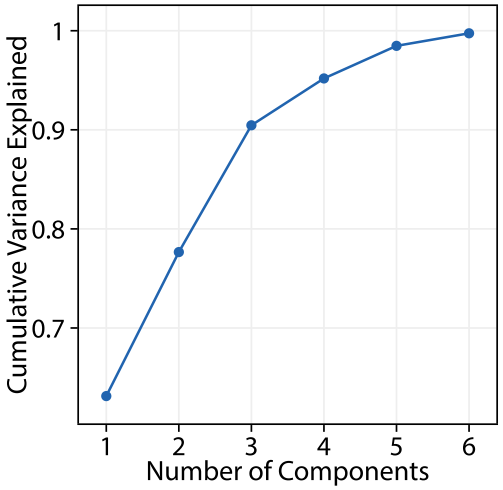


**Figure S5:** Variance explained by each principal component for principal component analysis performed on metabolite yields and growth rates of *E. coli* strains (described in Materials & Methods) grown in rich defined media in a bioreactor and microplates supplemented with reducing agents.

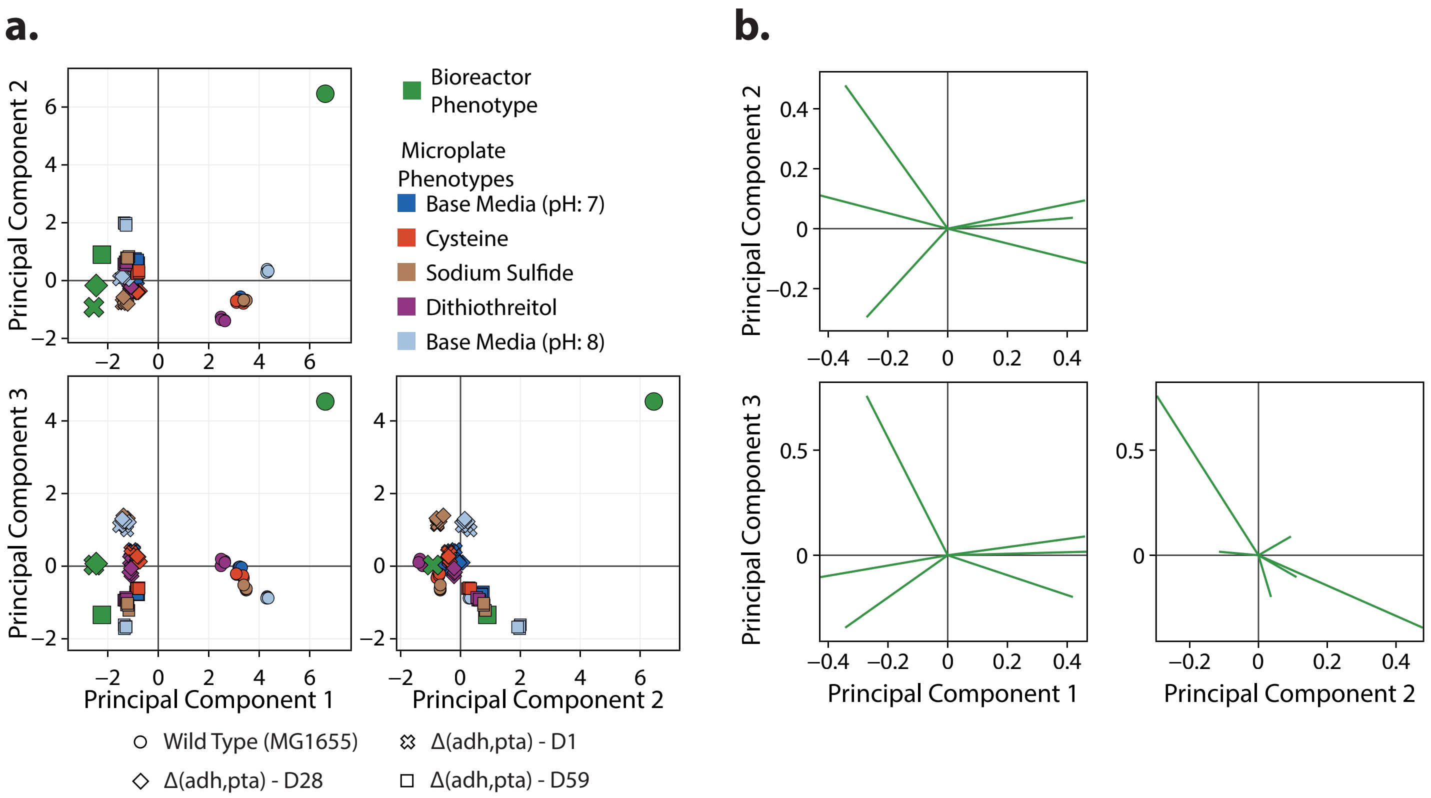


**Figure S6:** Principal component analysis performed on the metabolite yields and growth rates of *E. coli* strains (described in Materials & Methods) grown in rich defined media in a bioreactor and microplates supplemented with reducing agents. **a.** Scores and **b.** Loadings of each feature from PCA analysis.

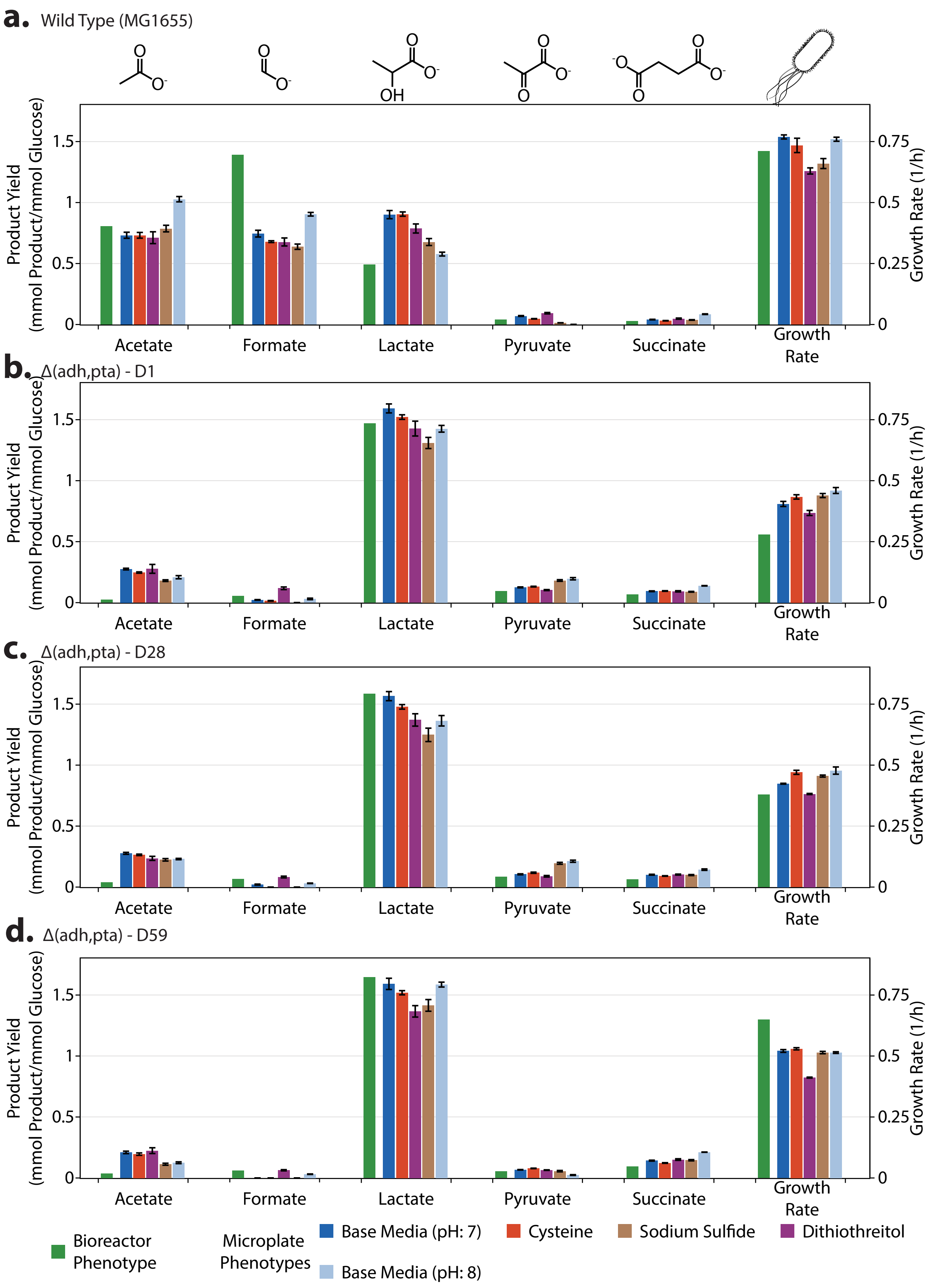


**Figure S7:** A comparison *E. coli*'s metabolite yields and growth rates obtained from a bench-top 0.5 L bioreactor and 96-well microplates with different reducing agents for the strains: **a.** Wild Type MG1655, **b.** MG1655 $\Delta(adhE, pta)$-D1, **c.** MG1655 $\Delta(adhE, pta)$-D28, and **d.** MG1655 $\Delta(adhE, pta)$-D59

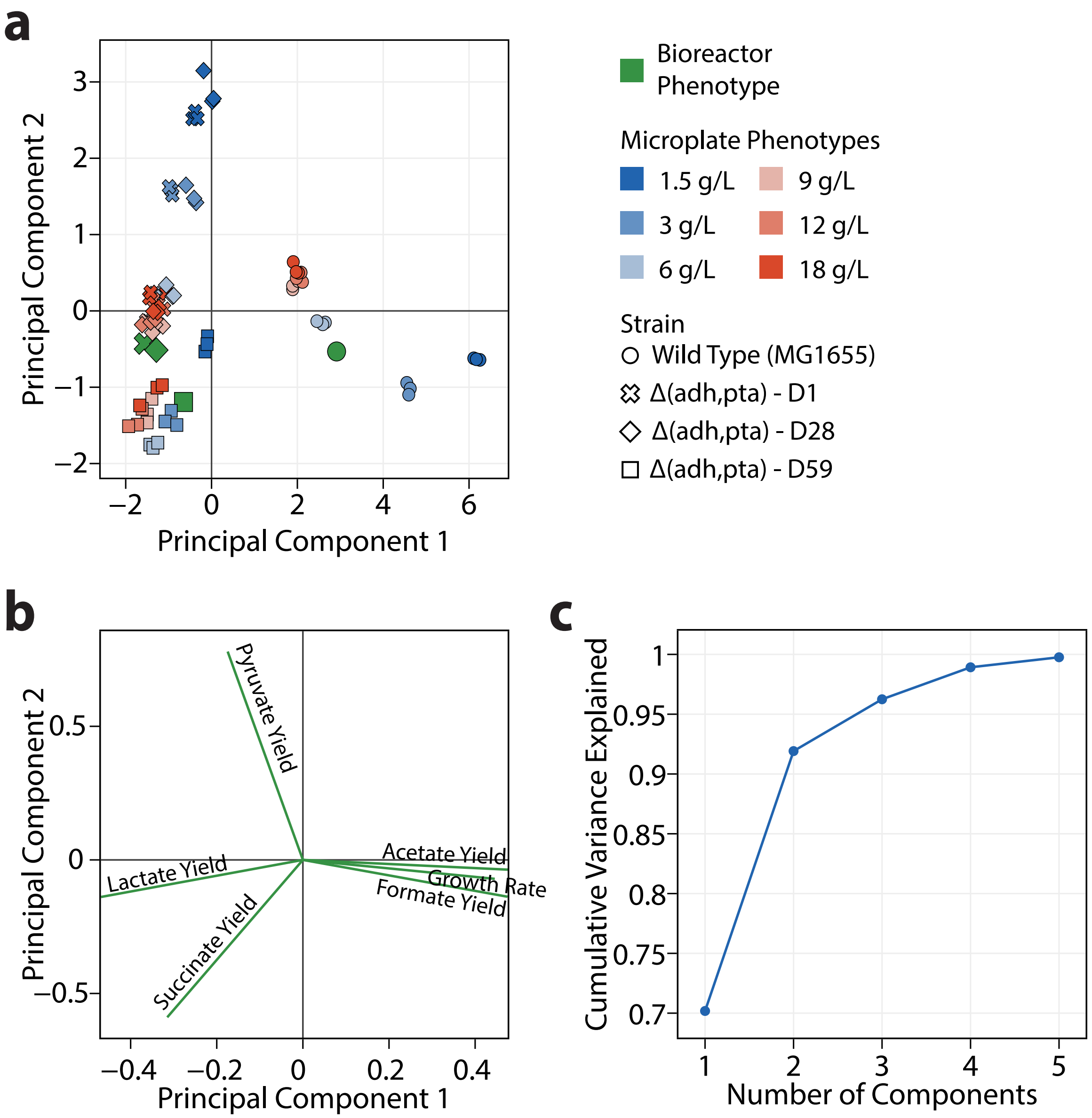


**Figure S8:** Principal component analysis performed on the metabolite yields and growth rates of *E. coli* strains (described in Materials & Methods) grown in rich defined media in a bioreactor and microplates supplemented with different substrate concentrations. **a.** Scores and **b.** Loadings of each feature from PCA analysis. **c.** Ratio of variance explained by each principal component.

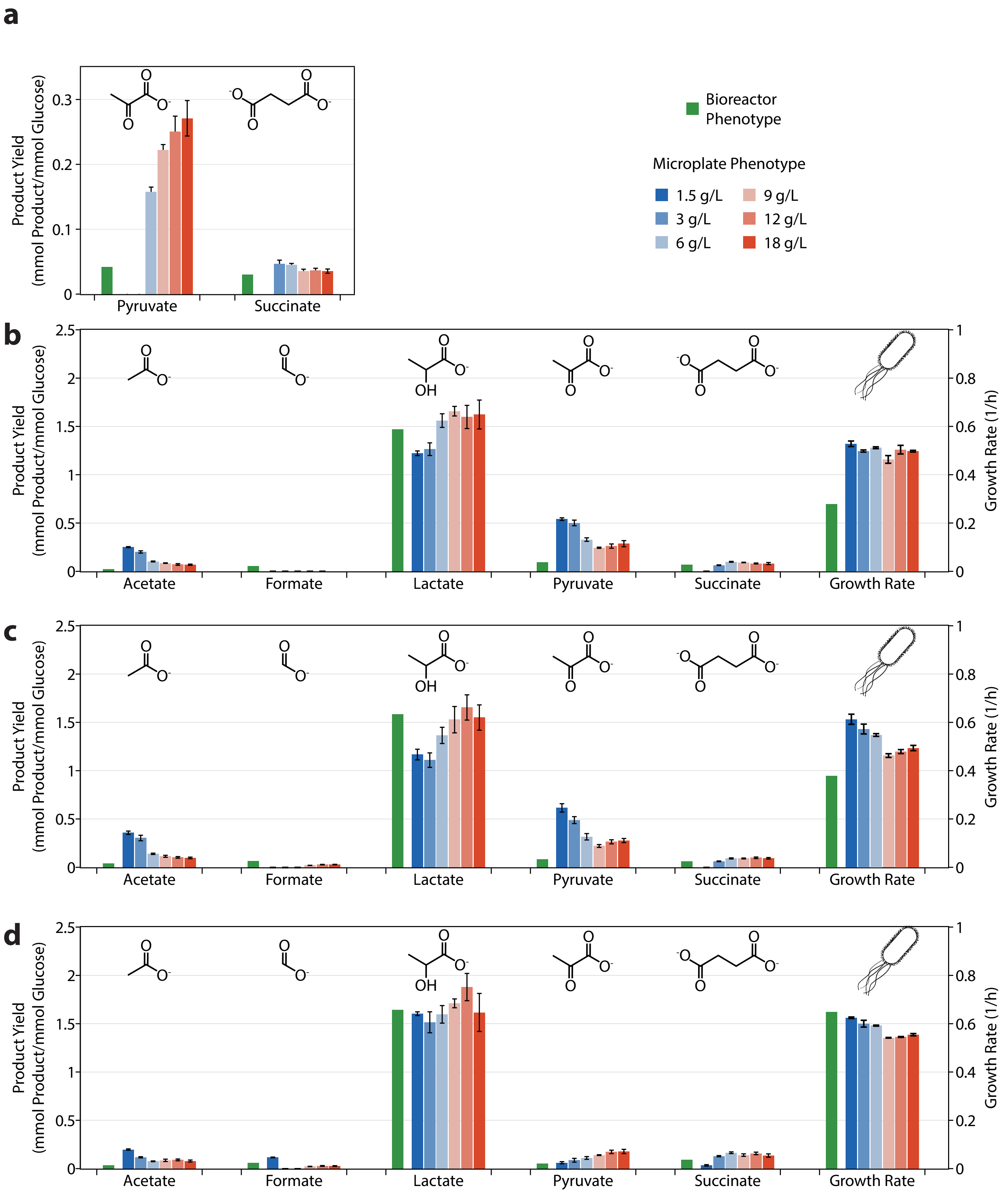


**Figure S9:** A comparison *E. coli*'s metabolite yields and growth rates obtained from a bench-top 0.5 L bioreactor and 96-well microplates with different initial glucose concentrations for the strains: **a.** Wild Type MG1655, **b.** MG1655 $\Delta(adhE, pta)$-D1, **c.** MG1655 $\Delta(adhE, pta)$-D28, and **d.** MG1655 $\Delta(adhE, pta)$-D59
